# Revealing the convergence of disease- and risk-associated circular RNAs in schizophrenia and bipolar disorder

**DOI:** 10.64898/2026.05.24.725548

**Authors:** Aarti Jajoo, Mateo Maya-Martinez, Nikolaos P. Daskalakis

## Abstract

Circular RNAs (circRNAs) remain an underexplored layer of transcriptomic regulation in psychiatric disorders. We quantified circRNA expression from 1,022 [518 neurotypical, 365 schizophrenia (SCZ) and 139 bipolar disorder (BIP)] postmortem cortex samples from PsychENCODE consortium cohorts and integrated these profiles with matched linear RNA and genotype profiles. We identified 23 SCZ-associated and 3 BIP-associated differentially expressed circRNAs (FDR<0.05; FDR-circDEG). We trained genetically regulated circRNA expression (circGReX) models using neurotypicals and applied them to SCZ and BIP GWAS to perform Transcriptomic Wide association analysis (TWAS) which identified 22 and 4 circGReX trait associations (circGTAs), respectively. Pathway enrichment of circDEGs and circGTAs implicated neuronal and synaptic processes for both disorders. In UK Biobank, circGReX–imaging associations were predominantly negatively correlated with SCZ and BIP circGTAs, but positively correlated with Alzheimer’s disease circGTAs. circKLHL24 isoforms showed the most prominent imaging associations. Many co-expression modules containing our FDR-circDEGs were enriched for psychiatric and neurodegenerative risk genes, including our identified circGTAs, and these modules were enriched for cognitive and neurodevelopmental traits. To conclude, circRNAs represent a distinct regulatory layer in psychiatric disorders, linking genetic risk to synaptic biology, brain structure and cognition through disease-specific expression, TWAS prioritization, and imaging associations.

## Introduction

Schizophrenia (SCZ) and Bipolar Disorder (BIP) are complex traits characterized by significant heritability, genetic overlap and a highly polygenic architecture involving numerous genes and biological pathways [1, 2]. The risk loci identified by the large-scale genome wide association studies (GWAS) [3, 4] typically fall within non-coding regulatory regions of the genome, influencing gene expression and splicing rather than directly altering protein structure or function. Genetic and brain transcriptomic data can be combined to understand the genetically regulated gene expression (GReX), which can be integrated with GWAS to conduct transcriptome-wide association studies (TWAS) [5, 6]. TWAS has advanced our understanding of the genetic basis of molecular mechanisms contributing to SCZ and BIP pathogenesis [7, 8]. Complementing these findings, molecular studies on postmortem brain tissues from subjects with SCZ and BIP have enabled us to distinguish a risk mechanisms from the ones downstream of disease pathology [9–12]. However, despite these efforts, we still do not fully comprehend molecular mechanisms of SCZ and BIP. Perhaps expanding beyond the gene-level linear RNA focus of existing studies could further improve our understanding. In fact, emerging studies of alternative splicing and other RNA families have already provided promising regulatory insights, underscoring the importance of broadening the molecular scope of psychiatric transcriptomic studies [12–15].

circRNAs are a special class of RNAs that have gained increasing attention over the last decade due to their unique circular structure and potential functional roles in various biological processes [16]. circRNAs are generated from similar precursor transcripts as linear RNAs, but through a distinct splicing process in which a downstream splice donor is joined to an upstream splice acceptor, forming a covalently closed loop [17]. Extensive research, particularly in the fields of cancer, cardiovascular, and immune-related diseases, has revealed diverse biological roles for circRNAs, including regulation of cell proliferation and differentiation, gene expression, and cellular stress responses [18]. Mechanistically, circRNAs function through various means, such as acting as microRNA (miRNA) sponges, binding proteins, modulating transcription of their host genes, and in some cases, being translated into peptides. Among many notable features, circRNAs are highly abundant in the brain [19, 20], where they are enriched in neuronal cell types and implicated in synaptic plasticity, and neurodevelopment. Their enrichment in the brain and potential involvement in neural processes have led to growing interest in circRNAs as contributors to psychiatric and neurodegenerative disorders.

Although research on circRNAs in psychiatric disorders is still in its early stages, several studies have begun to examine their expression in both brain tissue and peripheral blood [21–25]. To date, however, most psychiatric circRNA studies have been constrained by modest sample sizes, limiting statistical power and the ability to detect robust genome-wide significant associations. For example, differential expression analysis of circRNAs in cortical brain tissue from individuals with SCZ and Autism Spectrum Disorder (ASD) did not identify statistically significant results after multiple testing corrections [23, 25]. However, these studies demonstrated that circRNA expression is influenced by biological and genetic factors and identified putative miRNA-sponging–mediated downstream mRNA regulatory axes for potentially psychiatry-relevant circRNAs. Integration of circRNA expression with genetic variation to fine-map GWAS loci in Autism Spectrum Disorder and SCZ has also been explored, establishing potential circRNA-mediated trans-eQTL effects [24]. Additionally, studies in neurodegenerative disorder, particularly Alzheimer’s disease (AD) and Parkinson’s disease, have reported circRNAs with statistically significant differential expression after multiple-testing correction, underscoring the potential for circRNAs to capture disease-relevant biology even from standard ribosomal RNA–depleted RNA sequencing when adequately powered cohorts are available [26, 27]. Together, these observations highlight both the promise of circRNA profiling across brain disorders and a major gap in psychiatric research. In particular, large-scale multi-cohort studies integrating circRNA expression with genetic variation to improve statistical power, systematically comparing circular and corresponding linear RNA signals, linking circRNA variation to imaging and other population-scale phenotypes, and placing circRNA signals within broader molecular and functional contexts remain limited.

In this study, we leveraged ribosomal RNA-depleted DLPFC RNA-seq data from multiple PsychENCODE cohorts to construct a harmonized, circRNA-inclusive transcriptomic database of 1,022 unique postmortem cortical samples (Supplementary Table 1; Figure 1; Methods; [28]). This integrated dataset included 518 neurotypical controls, 365 SCZ subjects and 139 BIP subjects. We systematically tested for disease-specific associations of circRNA with SCZ and BIP. To uncover coordinated expression dynamics between circRNAs and linear RNAs, we constructed circRNA–mRNA co-expression modules using neurotypical control samples and then projected them to the SCZ and BIP data to identify the ones that were disease-associated. We then developed genetically regulated circRNA expression models (circGReX) using neurotypical control samples and applied them to GWAS summary statistics from SCZ and BIP, generating circRNA transcriptome-wide association study (circTWAS) results and identifying circRNA gene-trait associations (circGTAs). We also applied circGReX models to AD GWAS data, to compare disease specificity in the circGTAs across major psychiatric and neurodegenerative disorders. To assess broader endophenotypic relevance, we conducted a circGReX-based phenome-wide association study (circPheWAS) using UK Biobank brain imaging and behavioral phenotypes. We leveraged publicly available circRNA knowledge to contextualize the functional relevance of identified candidate circRNAs by constructing circRNA–miRNA–mRNA networks, checking for overlap with cell-type-specific expression profiles, conservation across species, evidence of translation, and functional roles supported by in vivo studies. Together, our framework identifies circRNA signatures of SCZ, BIP, and related endophenotypes, providing a foundation for follow-up translational circRNA-focused mechanistic studies.

**Figure 1.**
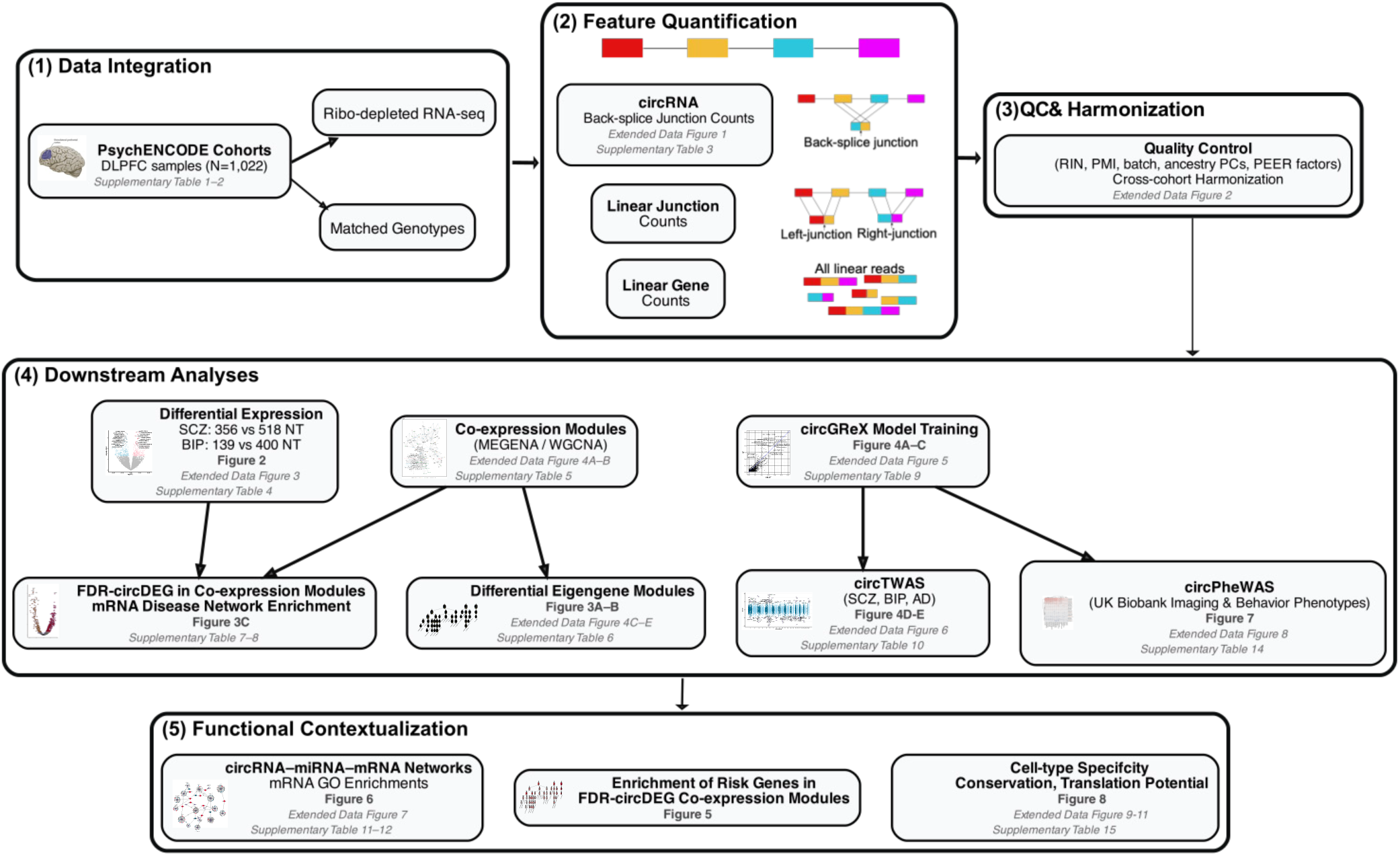
Overview of the circRNA-integrated transcriptomic analysis framework. Schematic overview of the analytical framework used to quantify and integrate circular RNA (circRNA), linear junction, and gene-level expression profiles across PsychENCODE postmortem cortex cohorts. (1) DLPFC ribo-depleted RNA-seq libraries and matched genotype data from PsychENCODE cohorts were integrated. (2) Feature quantification included circRNA back-splice junction counts, flanking linear junction counts, and linear gene-level counts. (3) Expression features were subjected to quality control and cross-cohort harmonization accounting for technical and biological covariates, including RNA integrity number (RIN), postmortem interval (PMI), batch effects, ancestry principal components (PCs), and PEER factors. (4) Downstream analyses included differential expression, co-expression network construction, circGReX model training, circTWAS, and circPheWAS analyses in UK Biobank imaging and behavioral phenotypes. Differential expression and co-expression analyses were further integrated to identify disease-associated modules and enriched gene networks. (5) Functional contextualization analyses included circRNA–miRNA–mRNA interaction networks, enrichment of psychiatric risk genes within co-expression modules, and characterization of cell-type specificity, conservation, and translational potential of prioritized circRNAs. Figure and supplementary references corresponding to each analysis step are indicated within the schematic.

## Results

### Multi-cohort integration and quantification of circular and linear RNA features in postmortem cortex

In our aggregated dataset, we quantified circRNA expression based on back-splice junction counts, together with linear RNA features, including gene-level counts and linear-junction counts corresponding to each circRNA (used as a proxy for the corresponding linear splicing event, defined as the mean of the left and right splice junctions flanking the circRNA) (Fig. 1A, Methods, Supplementary Note). After quality control, a total of 10,936 circRNAs were retained for downstream analyses (Methods, Supplementary Table 2). We refer to the host gene as the gene within which a circRNA is generated, defined by both back-splice junctions mapping to the same transcriptional unit; accordingly, circRNAs are labeled by their host gene using a standardized nomenclature that encodes transcript identity, strand orientation, and exon boundaries, with full details provided in the Methods and Supplementary Notes (Extended Data Fig. 1, Supplementary Table 3).

### Differential circRNA expression in SCZ and BIP reaches genome-wide FDR significance

RNA-seq libraries from multiple studies were analyzed with covariates selected to explain transcriptomic variance (Extended Data Fig. 2A-B, Methods). Our covariates did not correlate with disease status (Extended Data Fig. 2C-E), while they captured variations associated with technical variables, estimated cell type proportions and other expected confounders (Extended Data Fig. 2F-G). Differential expression analyses were performed on linear counts, and circRNA counts and their corresponding linear-junction counts (Supplementary Table 4; Methods), both at the individual-cohort level and using all the studies together as a mega-analysis. Mega-analysis results are presented as main, while cohort-specific results are provided in Supplementary Notes (Extended Data Figure 3). As a sanity check, we compared our linear differential expression results in SCZ and BIP with those reported by Gandal *et al.* [12]. Their study integrated RNA-seq datasets across the same cohorts used here, except that we excluded subjects that were profiles with poly(A)-selected libraries (26.6 % of the total subjects) because most circRNA lack poly(A) tails. Despite methodical differences in data processing and our exclusion of polyA-selected libraries, we observed strong concordance between z-scores of our nominal linear DE and those from Gandal et al, with correlations of 0.89 and 0.94 in SCZ and BIP, respectively (Methods, Extended Data Fig. 3A-F). Using a 5% false discovery rate (FDR) threshold, we identified 23 and 2 circRNAs that were significantly differentially expressed (FDR-circDEGs) in SCZ and BIP, respectively, (Fig. 2A-B, Method). This included *circHomer1* (Fig. 2C) which had been previously reported as downregulated in studies of psychiatric and neurodegenerative conditions [26, 29], but we demonstrated a genome wide FDR-significance for this gene in a psychiatric disorder for the first time (both in SCZ and BIP). Furthermore, we observed multiple FDR-significant circRNAs originating from a same host gene. Three downregulated circRNAs that derived from *ATRNL1* (2 in SCZ and 1 in BIP) and 2 downregulated circRNAs derived from *ATXN1* in SCZ (Fig. 2A-C). *circATRNL1* has also been reported to be downregulated at the FDR-significance level in a cortical AD study [26, 27].

**Figure 2.**
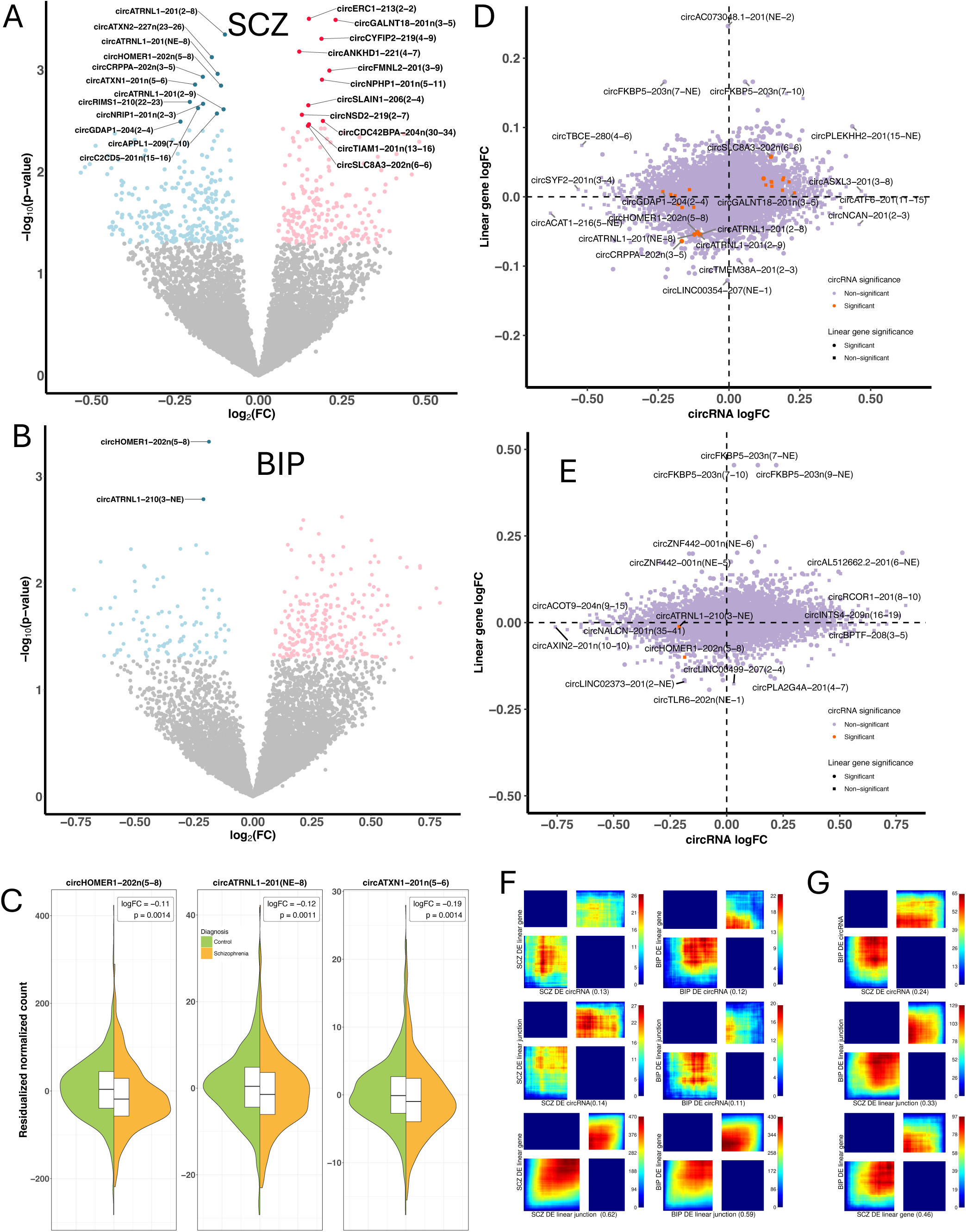
Gene-level circRNA differential expression, linear RNA concordance, and cross-disorder overlap in SCZ and BIP. **A. Volcano plot of differentially expressed circRNAs in SCZ**. Light colored dots represent nominally significant circRNAs (P < 0.05), with darker dots indicating those passing the FDR-adjusted threshold (P < 0.05). FDR-significant circRNAs are labeled. **B. Volcano plot of differentially expressed circRNAs in BP**. Light colored dots represent nominally significant circRNAs (P < 0.05), with darker dots indicating those passing the FDR-adjusted threshold (P < 0.05). FDR-significant circRNAs are labeled. **C. Residualized expression levels of top differentially expressed circRNAs between schizophrenia and control groups**. Violin and half-boxplots display normalized circRNA expression values after regressing out all differential expression analysis variables except diagnosis. Each subplot represents one circRNA. Annotated values indicate the log2 fold change and p-value from the differential expression analysis. **D-E. Comparison of logFC direction between circRNA and linear host gene RNA pairs in SCZ and BIP.** Shapes represent the statistical significance of the linear RNA (circles: significant; squares: non-significant), while colors indicate the significance of the circRNA (orange: significant; purple: non-significant). **F. Rank-rank hypergeometric overlap (RRHO) plots between differentially expressed circRNAs, linear genes and linear junctions within SCZ and BIP.** In this analysis, circRNAs, linear genes and junctions were ranked based on the product of the sign of log2(fold change) and −log10(p). The resulting RRHO heatmap illustrates the degree of overlap, with colors representing the −log10(p) values from the hypergeometric test assessing the significance of the gene list overlap. Warm colors in the bottom left and top right quadrants reflect overlap in genes with upregulation or downregulation, respectively, in both data sets. Warm colors in the top left and bottom right quadrants reflect overlaps in genes with opposite direction of effects in the two data sets. The Spearman correlation coefficient (ρ) based on the ranking metrics is provided as a reference for the effect size of the relationship. **G. Rank-rank hypergeometric overlap (RRHO) plots between differentially expressed circRNAs, linear genes and linear junctions across SCZ and BIP.**

### Weak correlation between circRNA and linear RNA disease associations

We observed a very low correlation between differential expression association effect sizes at the circRNA and linear-host gene levels (Pearson correlation of z-scores was 0.13 in SCZ and 0.12 in BIP; Fig. 2D-F). Similar low correlations were observed between effect sizes of circRNA and linear-flanking junctions (0.14 in SCZ and 0.16 in BIP; Fig. 2F). In contrast, the correlation between effect sizes of linear host-gene and linear flanking-junction counts associations were much higher (0.6 in SCZ and 0.58 in BIP). This suggests that the observed discordance reflects differences between circular and linear RNA disease signatures, rather than differences between junction-level and gene-level quantification. Rank–rank hypergeometric overlap (RRHO) analysis did not show enrichment in the upper-left and bottom-right quadrants, indicating an absence of systematic enrichment for opposite-direction log-fold changes (Fig. 2F). Nevertheless, approximately 36% of nominal circRNA–linear host-gene pairs exhibited opposite directions of effect. Cross disorder correlations of DE circRNA between SCZ and BIP were lower (0.24) than both linear-host (0.33) and linear-junction (0.46) signals (Fig. 2G).

### Co-expression networks include both circRNA–linear RNA modules and circRNA-only modules

We integrated circRNA and mRNA (protein-coding) to construct co-expression modules using MEGENA [30], restricting module construction to neurotypical control samples to capture correlation structures independent of disease processes (Supplementary Table 1). In total, we identified 846 hierarchical modules, of which 28% contained at least one circRNA (Supplementary Table 5; Extended Data Figure 4A-B). Although circRNA expression is known to decouple from their host gene expression [19], some degree of co-variation is expected due to shared genomic origin. Consistent with this, 19.3% of circRNAs co-membered with their host gene in at least one module. In addition, 21 modules contained only circRNAs. Together, these results indicate that circRNAs co-cluster with linear transcripts, including their host genes, while also forming circRNA-only co-expression modules.

### BIP-Associated Modules containing circRNAs are larger than SCZ-Associated Modules

To assess disorder-specific differences in coordinated co-expression, we tested MEGENA-derived modules for differential eigengene expression in SCZ and BIP using WGCNA (Methods; [31]). In total, 465 modules (321 upregulated) in SCZ and 393 modules (230 upregulated) in BIP were differentially expressed at FDR < 0.05 (Supplementary Table 6; Fig. 3A-B). Of these, 241 modules overlapped between the 2 disorders, this included 52 modules containing both circRNAs and linear mRNAs and none contained only circRNAs (Extended Data Fig. 4C-E). Module sizes of SCZ-specific modules containing both linear RNAs and circRNAs were significantly smaller than those observed in BIP (Wilcoxon test, p = 0.04; Extended Data Fig. 4D). Further, associations with circRNA-only modules were more frequent in BIP (n = 7) than in SCZ (n = 1).

**Figure 3.**
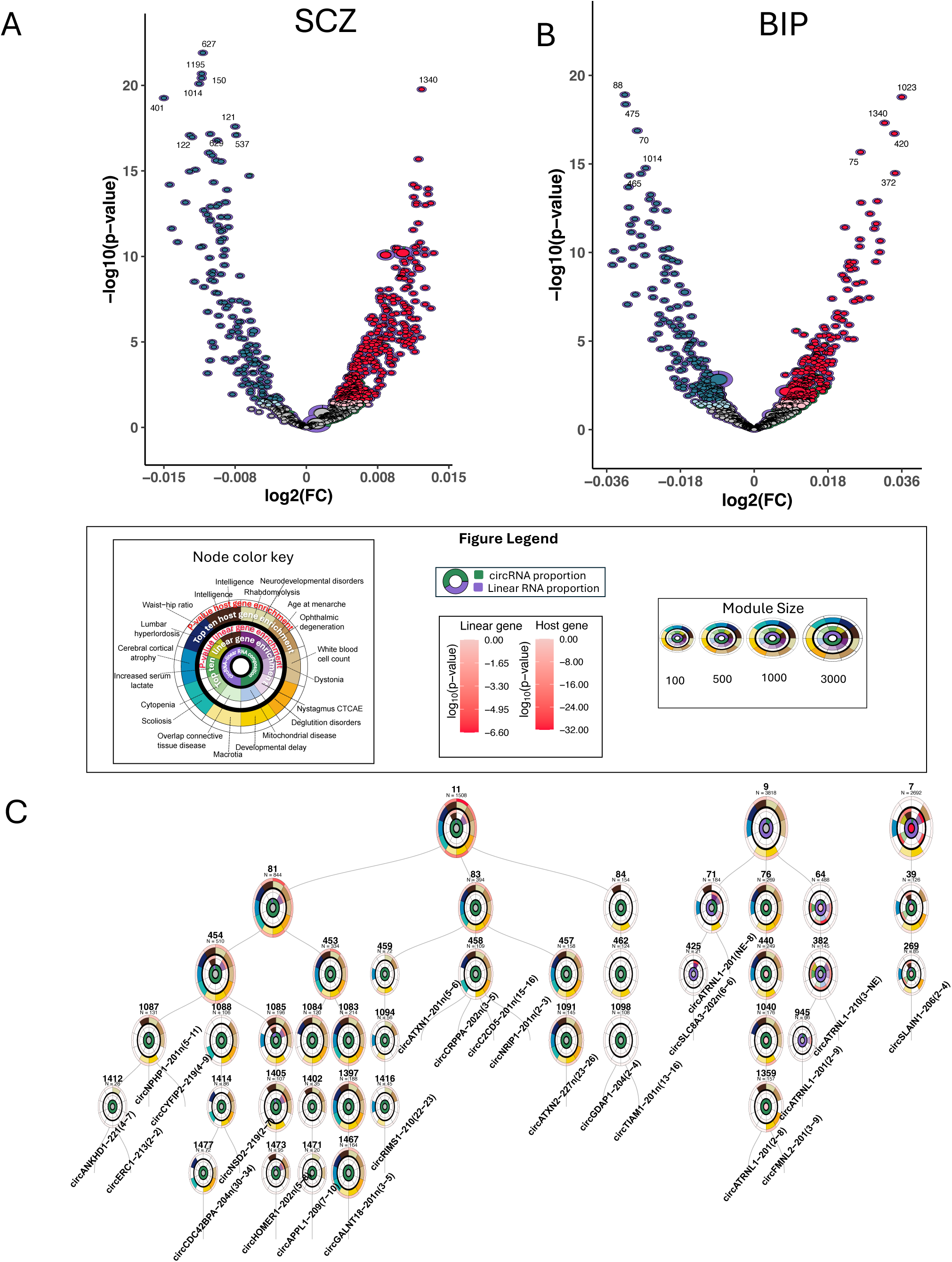
Differential module eigengene analysis and disease enrichment of circRNA-associated co-expression modules in SCZ and BIP. **A-B. Differential expression analysis of module eigengenes in SCZ and BIP.** Each circle is a MEGENA module. The ring around the circle shows the proportion of circRNA and linearRNA members in the module. Y-axis represent the differential eigengene logFC effect size, the analysis was performed using WGCNA package. **C. Hierarchical organization of co-expression modules harboring FDR-circDEGs in SCZ and BD, alongside DisGeNET disease enrichment profiles.** Each circular node represents a co-expression module, with branches illustrating parent–child relationships. Terminal child modules are annotated with the names of the FDR-circDEGs they contain. Within each module, the innermost circle represents the module eigengene differential expression logFC, with red and blue indicating up- and down-regulation, respectively, and color intensity reflecting statistical significance. The second ring shows the relative proportion of circRNA and linear RNA members within the module. The next two concentric rings summarize DisGeNET disease enrichments identified from linear RNA module members, where one ring denotes enriched disease categories and the adjacent ring indicates enrichment significance (-log₁₀(p-value)). The outermost two rings represent the corresponding DisGeNET disease enrichments and significance for circRNA host genes. Across all modules, the top 20 enriched disease categories are displayed, with colors in the disease enrichment rings corresponding to specific disease classes shown in the legend. Modules are labeled with module IDs and total member counts (N).

### FDR-circDEG-containing co-expression modules are enriched for neural and systemic disease phenotypes

The FDR-significant eigengene modules didn’t contain our FDR-circDEGs. We therefore focused on modules that contained FDR-circDEGs to explore disease relevance of their members. In total, 43 modules contained FDR-circDEGs, and were organized under three parent modules (M7, M9 and M11; Fig. 3C). M9 contains the largest number of FDR-circDEGs, and the majority of its members are circRNAs. Across FDR-circDEG–containing co-expression modules, both mRNA members and the host genes of circRNA-members showed enrichment for disease and trait phenotypes spanning neural, neurodevelopmental, and systemic domains (Supplementary table 7-8). Module 382 containing circATRNL1 FDR-circDEGs, was dominated by mRNA members. These mRNAs were enriched for cognitive, motor, mitochondrial and metabolic phenotypes, including intelligence, dystonia, mitochondrial disease, and serum lactate levels. Intelligence-related enrichment was observed across multiple modules at both the mRNA and circRNA host-gene levels, including module M1085, which also contained circHomer1. circHomer1 has been shown in animal models to regulate orbitofrontal cortex–mediated synaptic plasticity and cognitive behaviors [29]. Enrichment for age at menarche, a systemic endocrine developmental trait previously linked to schizophrenia onset [32], was observed across multiple FDR-circDEG–containing modules at the circRNA host-gene level.

### Heritable regulation of circRNAs and circGReX model development

We next quantified the extent of cis heritability regulation of circRNA expression and found the heritability estimates reaching up to 0.6 (Supplementary Table 9). The heritability correlation between circRNA and their linear host gene expression were low (r = 0.2; Fig. 4A). We trained three types of genetically regulated expression (GReX) models: circGReX (based on backsplicing counts), linear-junction GReX (based on linear-junction counts), and linear-gene GReX (based on linear gene level counts), using PrediXcan pipeline (Methods; [5]). We were able to predict 1359 circRNAs with cross-validated R² > 0.01 and p-value < 0.05, and these prediction R² values were positively correlated with circRNA heritability (pearsons’s correlation 0.84; Fig. 4B). For, 316 circGReX neither of the corresponding linear-gene or linear-junction level models were predictable. An additional 284 had corresponding linear-junction GReX predicted, and 243 had corresponding linearGReX predicted. Only 237 circRNAs had all three types successfully predicted (Extended Data Fig. 5A,B-D). Given that circRNAs and linear RNAs compete for splicing [33], we also expected SNPs to have opposite effect directions in the prediction models. Among the overlapped predicted genes, the SNP effect sizes between circGReX and linear-gene GReX models showed a relatively low correlation (r = 0.45), compared to a higher correlation between linear-junction GReX and linear-gene GReX models (r = 0.65) (Fig. 4C, Extended Data Fig. 5E-F). Several SNPs showed opposite direction of effect between circGReX and linear-gene GReX models, a pattern not observed between linear-gene and linear-junction GReX models confirming that observed differences reflect backsplicing vs linear splicing regulation (Extended Data Fig. 5E-F).

**Figure 4.**
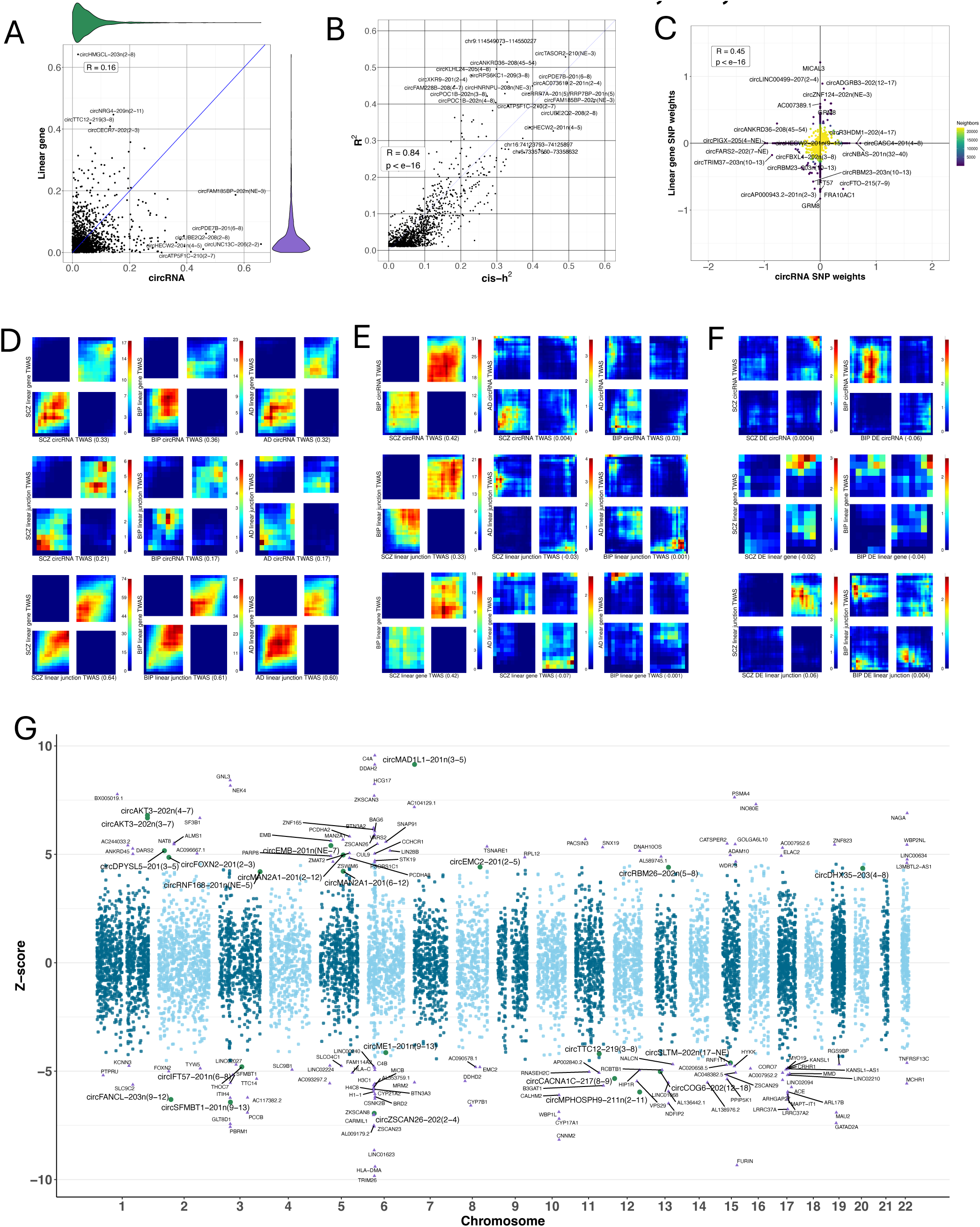
cis-heritable regulation, GReX modeling, and TWAS associations of circRNAs in SCZ, BIP, and AD. **A. Comparison of cis-heritability between circRNAs and their linear counterparts.** Scatterplot showing the cis-heritability (h²) of circRNAs (x-axis) versus their corresponding host gene’s linear RNA (y-axis). Marginal violin plots depict the distribution of h² values for each kind of RNA. **B. Relationship between cis-heritability and GReX model performance for circRNAs**. Scatterplot showing the correlation between circRNA cis-heritability estimates in the x-axis and cross-validated R² from GReX prediction models in the y-axis. **C. Comparison of SNP weights between circRNAs and linear gene.** SNP weights derived from GReX models trained on circRNAs (x-axis) and linear genes (y-axis) are shown. Points represent SNPs shared between models as well as SNPs present only in one model. Selected circRNAs and linear genes with extreme SNP weights across quadrants are highlighted. **D-F. Rank-Rank Hypergeometric Overlap (RRHO) analyses within and across disorders and analyses.** RRHO heatmaps depict the overlap of genes with positive and negative z-scores. Warm colors in the top-right and bottom-left quadrants indicate overlap of genes with the same direction of effect, whereas warm colors in the top-left and bottom-right quadrants indicate overlap of genes with opposite directions. Analyses were performed in three ways: within disorder (D), across disorders (E), and comparing differential expression with TWAS results (F). **G. SCZ TWAS results for circRNAs and linear genes.** TWAS results for circular RNAs (circles) and linear RNAs (triangles) associated with SCZ. Significant (Bonferroni-adjusted p-value < 0.05) circRNAs and linear genes are highlighted—green for circRNAs, purple for linear RNAs—and labeled.

### circTWAS showed low correlations with linear TWAS

We applied circGReX, linear-gene GReX, and linear-junction GReX to GWAS summary statistics of SCZ, BIP, and AD (Methods; Supplementary Table 10). To assess whether circGTAs capture distinct signals from their linear counterparts, we examined Spearman correlations of TWAS Z-scores and generated RRHO plots. As observed in DEGs, circGTAs showed low correlation with linear-junction GTAs (r = 0.20 for SCZ, 0.17 for BIP, and 0.17 for AD; Fig. 4D). The RRHO plots for AD showed enrichment in the second quadrant, indicating opposite Z-score directions for a significant subset of genes (Fig. 4D). circGTAs and linear-gene GTAs also showed modest correlation (0.30–0.36), whereas linear-junction GTAs and linear-gene GTAs were highly correlated (0.60-0.64) across all three traits. These patterns support that circGTAs capture distinct regulatory signals associated with disease risk.

### Expected cross-disorder circTWAS correlations and opposite TWAS-DE effects in BIP

Consistent with known genetic correlations [34, 35], we observed relatively higher GTA correlations between SCZ and BIP (r = 0.40 for circGTAs, 0.30 for linear-junction GTAs, and 0.40 for linear-gene GTAs), and much lower correlations with AD (r = –0.06 to 0.003) (Fig. 4E). When comparing SCZ and BIP to AD, RRHO plots revealed upper-left and bottom-right quadrant enrichments for linear-junction GTAs and linear-gene GTAs—indicating significant opposite effect directions. Notably, circGTAs showed relatively greater concordance, in comparison to linear-gene GTAs where the negative correlations observed. Finally, when comparing GTAs with DEGs for SCZ and BIP, we observed low overall correlation across features (Fig. 4F). Interestingly BIP comparison showed enrichment in the upper-left quadrant, indicating enrichment of downregulated circGTAs among upregulated circDEGs (Fig. 4F).

### circTWAS in SCZ, BIP, and AD reaches Bonferroni significance

Overall, we identified 22, 4, and 3 Bonferroni-significant circGTAs (bonf-circGTAs) for SCZ, BIP, and AD, respectively (Fig. 4G, Extended Data Fig. 6). Notably, *circSLTM* was a bonf-circGTA shared across all three traits with same direction of effect across the three disorders. *circSFMBT1* and *circMAD1L1* were bonf-circGTAs shared between SCZ and BIP, also showing same direction of effect. *circTTC3* was the only BIP-specific bonf-circGTA. Between SCZ and AD, *circZSCAN26* was a bonf-circGTA with opposite directions of effect, while *circCD2AP* was specific to AD. Within SCZ, *circFOXN2*, *circZSCAN26*, and *circEMC2* bonf-circGTAs exhibited effect directions opposite to those of their corresponding linearGTAs.

### FDR-circDEG co-expression modules enriched for TWAS signals

We found no overlap between FDR-circDEGs and bonf-circGTAs, a pattern also observed in linear transcriptomic analysis [36]. However, several co-expression modules containing FDR-circDEGs were enriched for our bonf-circGTAs and linear TWAS signals reported in TWAS atlas [37] across psychiatric and neurogenerative disorders (Fig. 5A; method). For example, M1473 which contains *circHomer1* was enriched for psychiatric TWAS risk genes (Fig. 3B). The module contains genes involved in synaptic signaling (*AKT3*, *UCHL3*), neuronal protein processing and quality control (*MAN1A2, NEMF*), and genome and RNA integrity (*APLF, CWC27, SDCCAG8*). Among these, *AKT3* and *SDCCAG8* have been implicated in SCZ TWAS [38], *UCHL3* in episodic memory TWAS [39], *MAN1A2* and *CWC27* single-cell TWAS of depression [40]. Another, example is M269, which contains FDR-circDEG *circSLAIN1* and shows enrichment for both bonf-circGTAs and linear SCZ TWAS signals. Risk genes in this module include *VRK2* and *FANCL* implicated in SCZ TWAS [7], *circMAN2A1* and *circFOXN2* are identified above as bonf-circGTAs, *MAN2A1* is implicated in both SCZ and AD [41]. *circMAN2A1* expression has also been associated with AD progression [42]. In addition, six circRNAs within M269 have been reported as differentially expressed in AD [26, 27] (*cirSLAIN1*, *circERBIN*, *circMAP7*, *circSTI8*, *circDOCK1* and *circMAN2A1*). Its parent module, M7, showed enrichment for AD risk gens. Other modules enriched for both psychiatry and neurodegenerative risk genes included M382 and its parent module M64.

**Figure 5.**
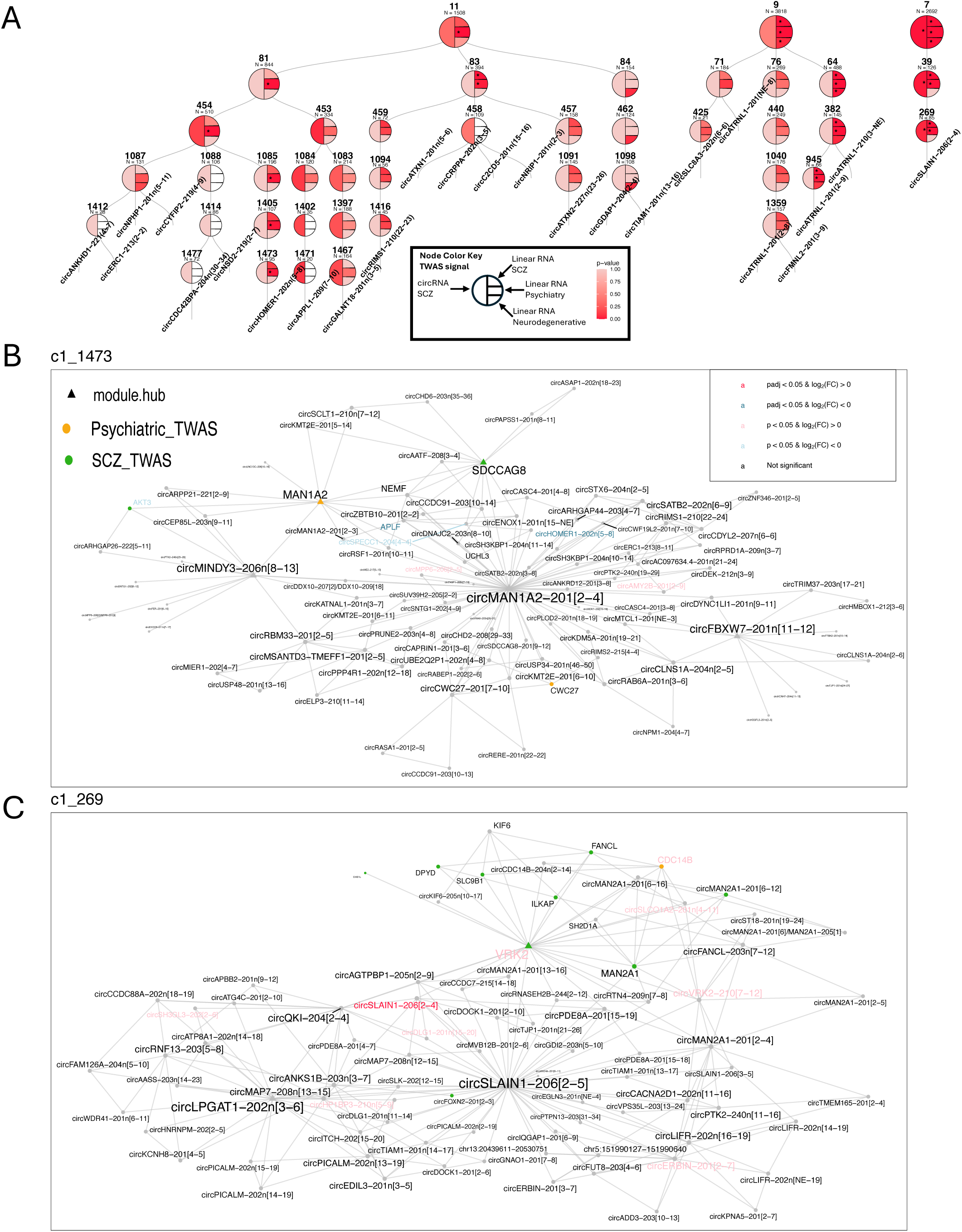
Convergence of TWAS-prioritized risk genes within FDR-circDEG co-expression modules. **A. Hierarchical organization of co-expression modules showing different TWAS enrichments patterns.** The plot follows the same hierarchical structure as Figure 3C. Nodes represent co-expression modules containing FDR-circDEGs, where parent-child relationships reflect hierarchical module organization. Bottom-most nodes are labeled by the corresponding FDR-circRNA members contained within each module. Each node is partitioned into four sections reflecting enrichment significance of TWAS-prioritized genes among module members: SCZ circRNA TWAS (left half), SCZ linear TWAS (upper-right quadrant), psychiatric disorder linear TWAS (middle-right quadrant), and neurodegenerative disorder linear TWAS (lower-right quadrant). Color intensity corresponds to enrichment p-values from Fisher’s exact tests, and * indicates p-value < 0.05. **B-C.** Examples of co-expression modules containing risk genes. **5B**. Co-expression network of module M1473 containing circHomer1. Nodes represent genes and circRNAs within the MEGENA module, while edges indicate inferred co-expression relationships between connected features. Triangles denote hub genes identified based on network connectivity. Node colors indicate TWAS-prioritized genes, with orange representing psychiatric disorder TWAS associations and green representing SCZ TWAS associations. Text colors indicate differential expression results: black for non-significant features, pink/red for positive logFC, and blue for negative logFC, with darker shades representing stronger statistical significance. **5C** Co-expression network of module M269 containing circSLAIN1, shows using same annotation schems as in panel B.

### circRNA–miRNA–mRNA networks link FDR-circDEGs and bonf-circGTAs to shared and distinct downstream mRNA programs

One mechanistic means by which circRNA regulates mRNA is through miRNA sponging [43]. We bioinformatically identified downstream mRNA targets of our FDR-circDEGs and bonf-circGTAs, and obtained circRNA-miRNA-mRNA axis using CirNetVis (Extended Data Figure 7A-B; Supplementary Table 11-12, Methods; [44]). Gene ontology enrichment of mRNAs in circRNA–miRNA–mRNA axis revealed both shared and distinct functional programs between FDR-circDEGs and bonf-circGTAs (Fig. 6A-B). Across both analyses, enriched terms included canonical Wnt signaling and broad neuronal projection/differentiation programs (Fig. 6A–B). In FDR-circDEG networks, enrichment was additionally dominated by chromatin and nucleosome organization, chromosome/telomere-related processes, and morphogenesis-related developmental programs (Fig. 6A, Extended Data Figure 7C). In contrast, bonf-circGTA networks showed stronger enrichment for TGF-β–related responses, receptor activity and phosphorylation regulation, and vesicle-mediated trafficking/secretion processes (Fig. 6B, Extended Data Figure 7D).

**Figure 6.**
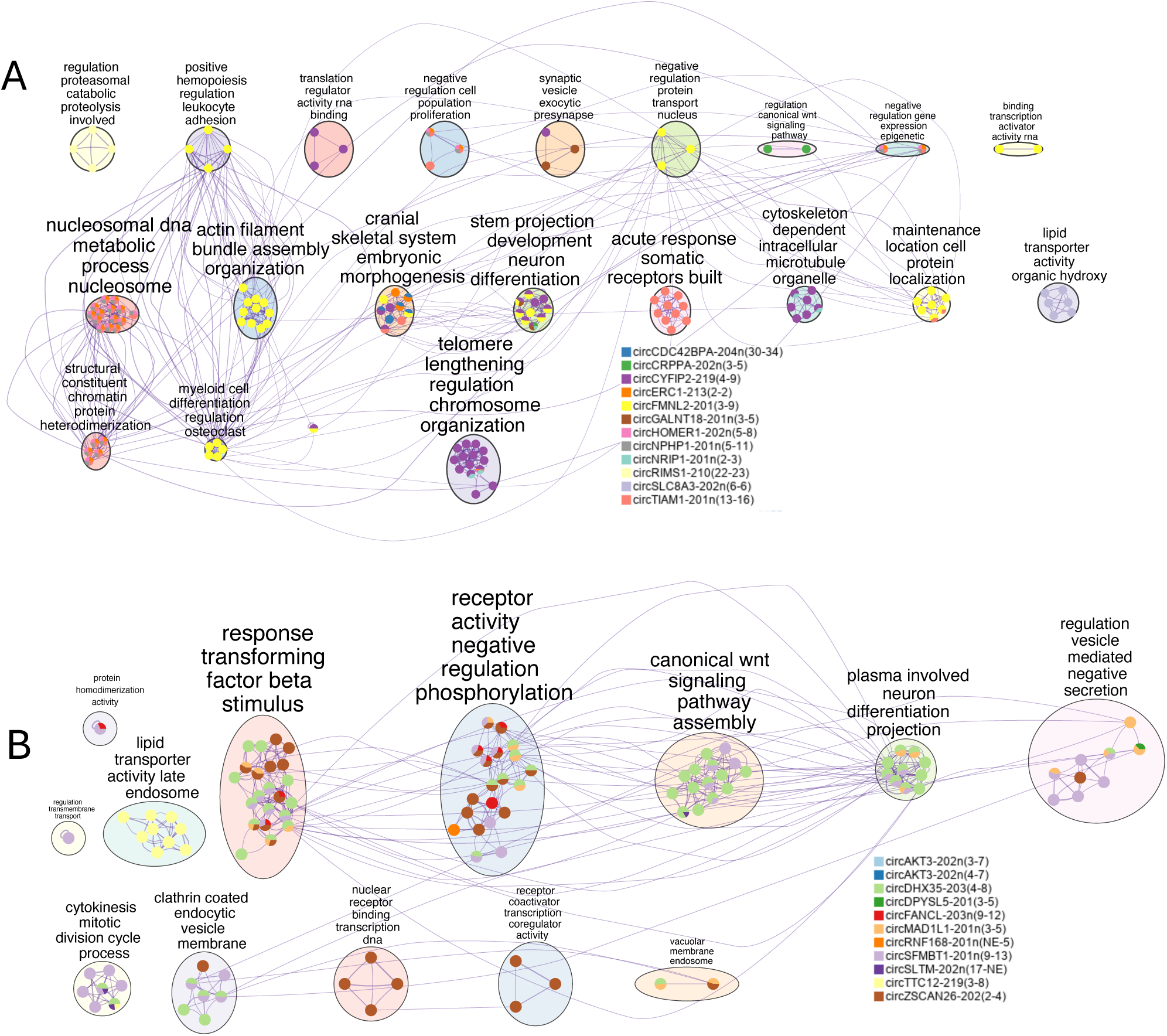
circRNA–miRNA–mRNA interaction networks reveal enriched biological processes associated with FDR-circDEGs and bonf-circGTAs. Metascape enrichment network plots showing biological processes associated with predicted circRNA–miRNA–mRNA interaction networks derived from FDR-circDEGs (**A**) and bonf-circGTAs (**B**). Nodes represent enriched biological process terms, while edges connect functionally related terms based on similarity. Pie charts indicate the relative contribution of interaction networks from individual circRNAs, highlighting both shared and circRNA-specific functional enrichments.

### Widespread circRNA associations with neurite density measures

Using circGReX models, we imputed circRNA expression in the UK Biobank cohort subjects and tested associations with observed brain imaging phenotypes (Methods, Supplementary Table 13). We focused on one of the most heritable classes of imaging-derived phenotypes, comprising 48 intracellular volume fraction (ICVF) measures that index neurite density [45]. Genetic correlations between ICVF measures and SCZ or BIP were low and positive, whereas correlations with AD were negative. In contrast, circGReX–ICVF association z-scores derived from SCZ and BIP circTWAS were predominantly negative, and associations observed using AD GWAS showed the same direction but substantially smaller magnitudes (Fig. 7A). Across these IDPs, we identified between 1 to 13 Bonferroni-significant circGReX (bonf-circGReX) per ICVF variable, corresponding to a total of 42 distinct bonf-circGReX. Several circRNAs were associated with multiple IDPs, with the strongest signal observed at the *KLHL24* locus (Fig. 7B, Extended Data Fig. 8A). This locus harbored nine Bonferroni-significant circGReX isoforms, with individual circRNA isoforms associated with up to 32 IDPs. We also observed multiple mitochondrial and oxidative stress–related genes within module M71, which contains many circKLHL24. This included circSLC8A1 (also upregulated in SCZ), which has been shown to be upregulated under oxidative stress in cell culture experiments and has been implicated in stress-related Parkinsonism [46]. *circKLHL24* has also been reported to regulate the miR-1204/ALX4 axis [47]. *ALX4* plays a role in skull and craniofacial development and shows higher expression during brain development [48]. Using *circKLHL24-*centered circRNA-miRNA-mRNA axis from Chen et *al.* [25] which was derived from matched circRNA, mRNA and miRNA data rather than sequencing-based predictions alone, we observed enrichment for multiple cognitive and neurodevelopmental traits (Fig. 7D). Additionally, other bonf-circGReX associated with multiple ICVF measures included *circNMT1* (19 IDPs), *circATF1* (13 IDPs), and *circZSCAN26* (10 IDPs). We also found that 4 of our SCZ bonf-circGTAs, *circAKT3* (2 isoforms), *circDPYSL5*, and *circZSCAN26* (also bonf-circGTA in AD, albeit with opposite direction of associations), were associated with many ICVF measures. *circANKHD1*, identified as our SCZ FDR-circDEG, was also a bonf-circGReX.

**Figure 7.**
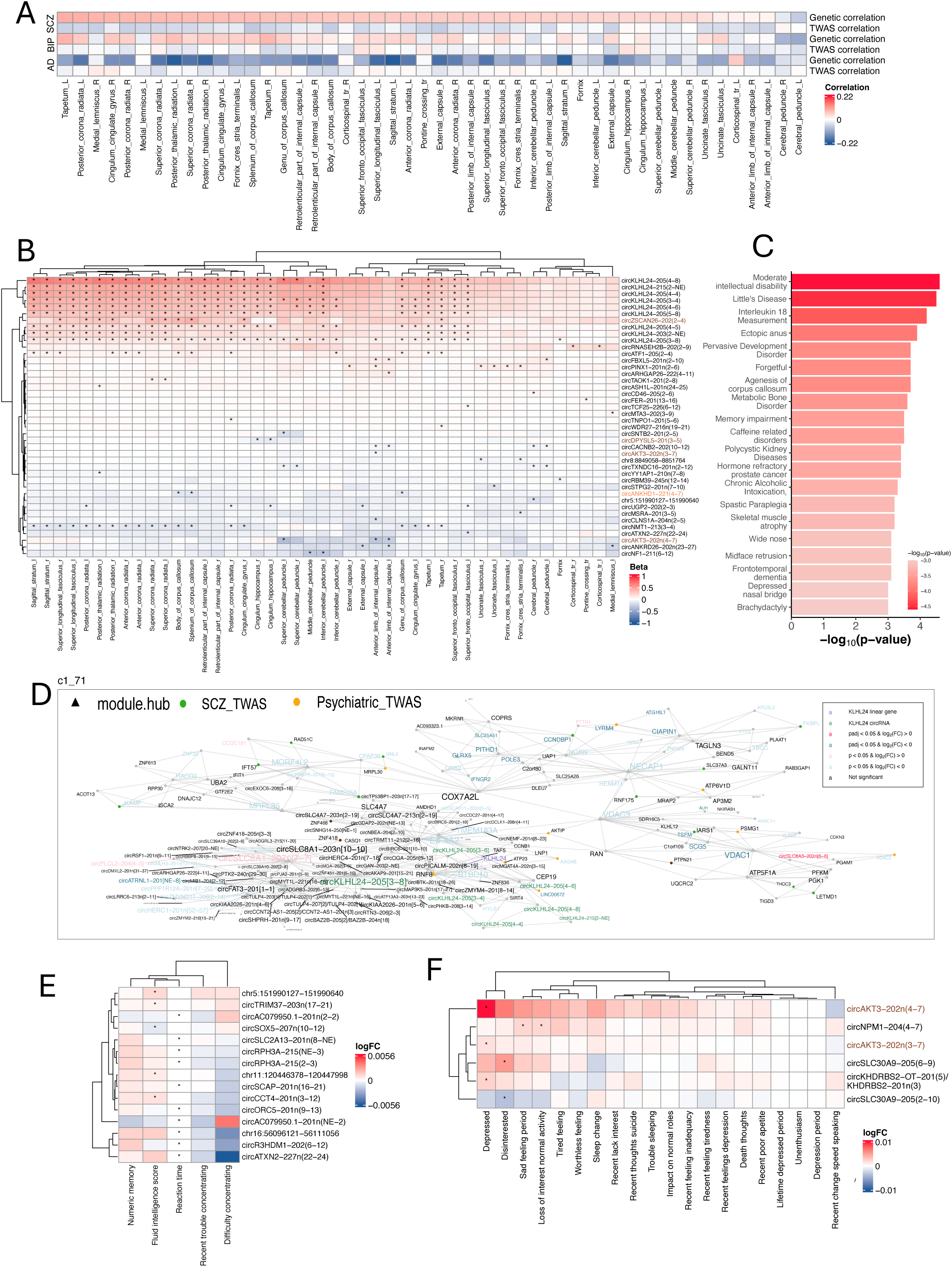
circGReX-Imaging & circGReX-Phenotype. **A. GWAS-based genetic correlations and circTWAS-based correlations between disorders and ICVF variables.** Heatmap showing GWAS-based genetic correlations between SCZ, BIP, and AD and ICVF imaging-derived phenotypes, together with TWAS based correlations between circTWAS disorder signals and circGReX-imaging association profiles across ICVF variables. Color intensity reflects the direction and magnitude of correlations. **B. circGReX-ICVF association based ICVF variable clustering** Heatmap showing clustering of ICVF imaging-derived phenotypes based on circGReX–ICVF association profiles. Each cell is colored by the z-score direction of the circGReX–ICVF association test, with red indicating positive effects and blue indicating negative effects. Significant associations are marked with * at the center of each cell. Names of FDR-circDEGs and bonf-circGTAs are highlighted in orange and dark brown, respectively. **C. Pathway enrichments of downstream genes associated with circKLHL24 through the circRNA–miRNA–mRNA interaction network.** **D. Co-expression module containing circKLHL24.** The plot follows the same structure as Figures 5A and 5B. **E-F. Associations between circRNAs and UK Biobank phenotypes.** Each heatmap displays the direction of circRNA associations—red for positive effects and blue for negative—across cognitive (**E**) and behavioral (**F**) phenotypes. Significant associations are marked with a star at the center of each cell.

### circGReX associations with cognitive and behavioral phenotypes

We also associated five cognitive variables with imputed circGReX in UK Biobank and found that two—fluid intelligence score and reaction time—were Bonferroni significant, with five and ten non-overlapping circRNAs, respectively (Supplementary Table 15). Among these, another circATXN2 isoform was also identified as a FDR-circDEG. In addition, we tested 20 behavioral variables and found that four showed Bonferroni-significant associations, involving six circRNAs in total. Among these were 2 isoforms of circAKT3, both of which were Bonferroni-associated with several ICVF measures and are also among our bonf-circGTAs in SCZ. (Fig. 7E-F, Extended Data Fig. 8B).

### circRNA enriched for pyramidal neurons, and show variable level of conservation and protein translation potential

To further contextualize the functional relevance of our circRNA findings, we integrated multiple layers of external circRNA annotations (Fig. 8, Extended Data Figure 9-11). First, we assessed evolutionary conservation using the Multiple Conservation Score (MCS) [49], which aggregates data from six species across 19 tissues. Among our detected circRNAs, 95% were found in the MCS database, with an average conservation score of 3.6. MCS values for FDR-significant circDEGs, bonf-circGTAs, and bonf-circPheWAS circRNAs spanned a broad range (2.0–6.5), indicating variable evolutionary constraints. Notably, *circHOMER1* showed a moderate conservation score, while *circKLHL24* was among the highest scoring (MCS = 6.4). Other highly conserved circRNAs included *circFMNL2* (MCS = 7.0), *circRIMS1*, and *circAKT3*.

**Figure 8.**
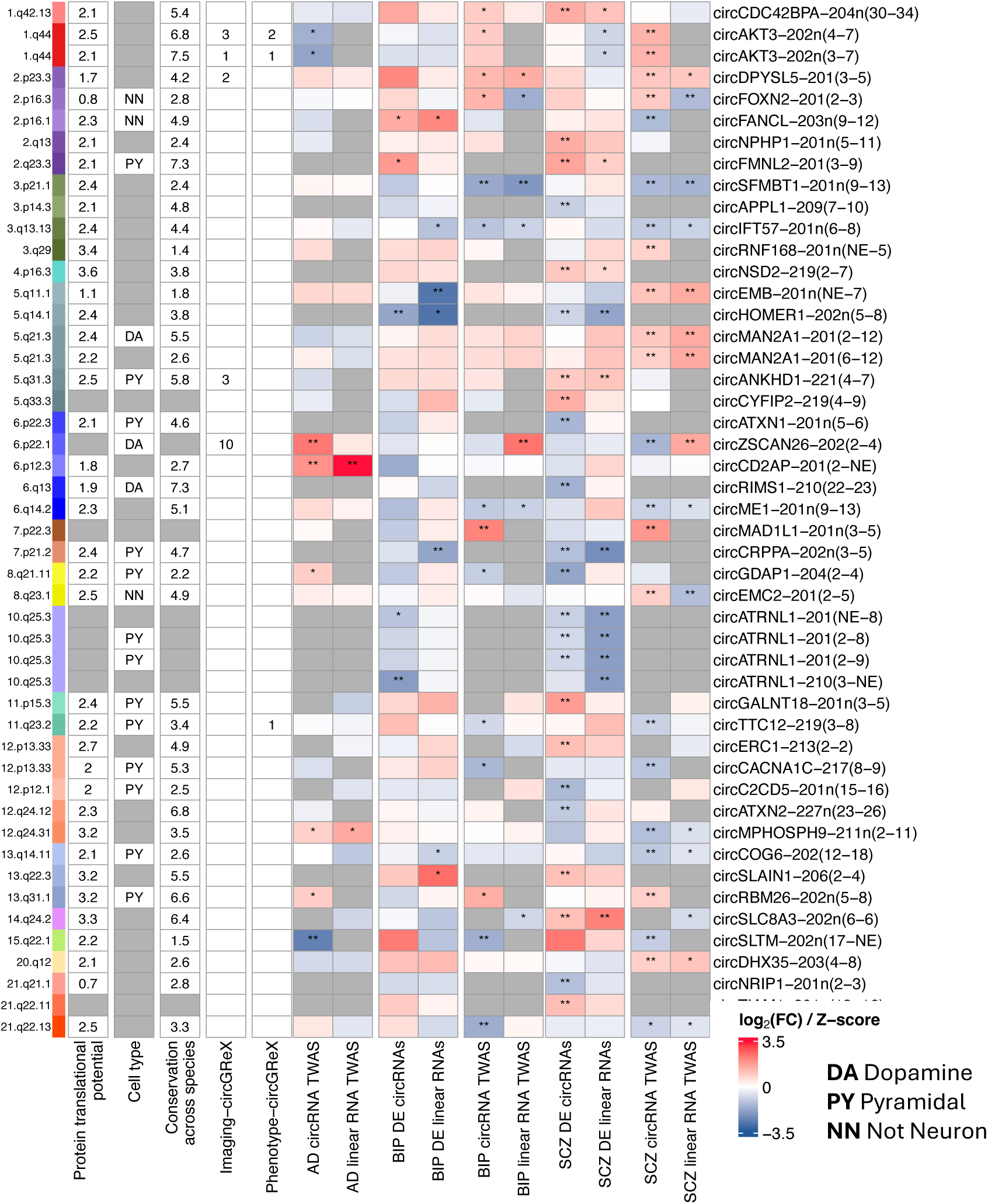
Summary of FDR-circDEGs and bonf-circGTAs identified in SCZ and BIP based on differential expression, TWAS and UKBB associations analysis, and existing annotation. Rows represent circRNAs identified as FDR-circDEGs or bonf-circGTAs in SCZ and BIP analyses. Columns summarize differential expression and circRNA TWAS signatures across SCZ, BIP, and AD, together with circGReX associations with UKBB imaging-derived phenotypes and additional circRNA annotations, including protein translation potential, cell-type specificity and conservation across species. Heatmap cells are colored by effect direction and magnitude, with red indicating positive associations and blue indicating negative associations. Significant associations are marked with * at the center of each cell.

Next, we examined cell-type specificity using annotations from a study profiling circRNAs in dopamine and pyramidal neurons (neuronal), and plasma fibroblasts (non-neuronal) [50]. Among the 10,000 circRNAs in our dataset, 529 were dopamine-specific, 1,457 pyramidal neuron-specific, and 1,653 non-neuronal. Strikingly, all FDR-significant SCZ circDEGs were classified as neuronal, with the majority being pyramidal-specific, showing significant enrichment (Fisher’s exact test, p = 0.00167; odds ratio = 12.54). However, gene set enrichment analysis (GSEA) of pyramidal-specific circRNAs showed only weak enrichment, suggesting that SCZ-associated circRNA dysregulation is driven by a subset of pyramidal-neuron circRNAs rather than a broad perturbation across all neuronal circRNAs. One of the top circGReX-Imaging hits, *circKLHL24*, was classified as pyramidal neuron-specific as well. Next, we assessed translational potential using the TransCirc database [51], which integrates evidence from m⁶A modifications, open reading frames (ORFs), and ribosome binding data. While most circRNAs had modest translatability scores, a subset—including *circNSD2*, *circRNF168*, and *circUGP2*—were predicted to have high (>3) translational potential. *circKLHL24* consistently scored highly across all categories, supporting its potential to be translated into a functional peptide. And *circMAN2A1* is already shown to be translated into a peptide [42].

## Discussion

By integrating postmortem cortex transcriptomic and genetic data across multiple PsychENCODE cohorts, we identified genome-wide significant circDEGs and circTWAS-prioritized circGTAs in both SCZ and BIP. We observed limited correlation between circular and linear RNA signatures across both differential expression and TWAS analyses, suggesting circRNAs capture disease- and disease-risk associated biology is not fully captured by canonical linear transcriptomic measurements and analyses. The identification of previously studied circRNAs, including circHomer1, circSLAIN1 and circMAN2A1, despite the limited number of existing neuropsychiatric circRNA studies, increases confidence that the numerous additional circRNAs identified in our study represent plausible disorder-associated candidates, substantially broadening the circRNA landscape implicated in these disorders. Although the functional roles of circRNAs remain incompletely understood, contextualization of our identified circRNAs using existing datasets provided important biological insight. We found that FDR-circDEGs and bonf-circGTAs were enriched in pyramidal neurons, emphasizing cell-type specificity. Modules containing FDR-circDEGs included numerous genes with known neuronal functions and were enriched for TWAS-prioritized psychiatric risk genes, suggesting network-level convergence between disease-associated circRNAs and genetic risk. Shared enrichments for neuronal differentiation and signaling pathways alongside distinct chromatin- and TGF-β–related programs among putative downstream mRNA targets inferred through miRNA sponging suggest that circRNAs may contribute to psychiatric disease biology through multiple regulatory mechanisms. Finally, strong associations between circGReX and neurite-density imaging traits, including widespread associations involving circKLHL24, suggest a mechanistic link between circRNA regulation and brain microstructure. The predominantly anticorrelated relationship between imaging-associated circGReX signals and SCZ/BIP circTWAS signals further suggests that psychiatric genetic risk may may result in reduced neurite density through circRNA-mediated regulatory mechanisms.

circRNAs and their host gene linear transcripts arise from the same genomic loci [52]. But prior studies support independent variation of circRNA and linear RNA expression during response to synaptic plasticity and neuronal differentiation, and across diverse cell types [19, 20]. Consistent with these observations, we also find substantial divergence between circRNAs and their host linear RNAs in our analyses. Notably, correlations between circRNA and linear RNA effects are higher in TWAS than in differential expression analyses, which may reflect disease-related processes that precipitate over the lifespan divergence between circRNA and linear RNA regulation. Approximately 36–40% of nominal circRNA–linear RNA pairs showed opposite directions across differential expression and TWAS analyses, which may reflect competitive regulatory relationships between circRNAs and their host linear RNAs [33, 53]. For example, circSLTM, one of the few bonf-circGTAs shared across BIP, SCZ, and AD, showed an association opposite in direction to a linear SLTM isoform [7] despite the absence of association at the gene-level count. Notably, this locus has also been reported to show shared colocalized loci for AD and SCZ specifically in putamen tissue, among six brain tissues tested in that study, including cortex [54]. This highlight both splicing- and cell-type–specific genetic regulation. Cross-order differential expression correlations show higher degree of separation in circRNA signatures compared to linear RNA between BIP and SCZ. Also, downstream mRNA signals linked to circRNA associations and circRNA-containing co-expression modules showed limited enrichment for inflammatory pathways, indicating that circRNA signatures capture molecular aspects of SCZ and BIP that are distinct from canonical inflammation-related processes. Together, these findings suggest that circRNAs provide complementary biological information beyond linear RNA, helping to fill an important gap in our understanding of psychiatric disease biology and offering a potentially informative repertoire for biomarker development.

Despite the decoupled circRNA and host linear RNA expression, shared regulation still produces pockets of coordinated expression. The presence of direct edges linking circRNAs to their host genes in several of our MEGENA modules captures this co-expression and reinforces the robustness of our modules. In general, these co-expression modules capture coordinated regulation but not causal relationships. The mRNA and circRNA members within a module may reflect mRNAs acting upstream of circRNAs, downstream of them, or both being jointly regulated by latent factors. We also observed several co-expression modules composed exclusively of circRNAs, and several FDR-circDEGs were contained within these modules. One possible explanation is that back-splicing is influenced by regulatory factors such as transcription or splicing regulators that are not captured within the modules, potentially because their effects reflect local availability or activity rather than global expression variation. Further work will be required to delineate the upstream mechanisms governing circRNA regulation and to clarify the downstream functional roles of these circRNA-only modules.

Risk gene enriched co-expression modules have been identified in the past [55, 56], and interestingly several of our modules containing FDR-circDEGs were enriched for psychiatric risk genes. For example, the *circHomer1*-containing module was enriched for psychiatric risk genes. In addition, genes within the module were enriched for cognitive traits, including intelligence, aligning well with prior studies linking circHomer1 to synaptic plasticity and cognition [29]. Another example *circSLAIN1-*containing modules that show enrichment for TWAS prioritized linear genes in SCZ and AD, TWAS prioritized circRNA in SCZ, and several differentially expressed circRNAs in AD. The convergence of these observed disease processes and risk signals across both linear and circular RNA within a single module suggests that it may represent a crucial molecular network for pleiotropic brain functions positioned at the interface of neuropsychiatric and neurodegenerative disease biology. Many risk genes within these modules are not differentially expressed, may imply subtle or synergistic contributions that act downstream rather than through strong individual expression changes. Alternatively, these genes may exert critical effects during neurodevelopment, with their influence persisting into adulthood through circuit-level vulnerabilities. Disease progression and chronic pharmacological exposure may further reshape transcriptomic landscapes, creating a gap between genetic risk signals and observed disease-state expression. In BIP, larger circRNA modules with differential eigengenes suggest synchronized expression patterns, possibly reflecting homogeneous drug response under monotherapy [57]. Supporting this interpretation, we observed enrichment of downregulated circTWAS associations among downregulated differentially expressed circRNAs in BIP, raising the possibility that some disease-associated circRNA signatures may reflect pharmacological attenuation of genetically driven expression patterns. Together, these findings underscore the dynamic interplay between genetic risk, disease pathophysiology, and treatment effects, highlighting the potential of circRNA-inclusive molecular networks to provide a more complete view of psychiatric disease biology.

Association between circGReX and diffusion MRI-derived ICVF variables may be biologically plausible given that circRNAs are markedly enriched in neuronal processes and synapse-associated compartments [19, 20, 58], including dendritic, axonal, and synaptic terminals, which represent the same microstructural densities captured by ICVF variables. Interestingly, while genome-wide genetic correlation between neurite density and SCZ is positive, circRNA TWAS signals show a negative correlation, which could reflect technical or global aggregation limitations of genetic correlations. While little is known about circKLHL24, the convergence of imaging associations, co-expression network structure, and downstream regulatory relationships surrounding this locus suggests that this locus may reflect broader neurodevelopmental and neurobiological processes relevant to psychiatric and neurodegenerative disease.

Until recently, circRNA annotations remained incomplete, and most back-spliced junctions were identified through de novo discovery approaches. Today, nearly all circRNAs identified in our study are already annotated, yet their functional roles remain largely unknown. There are no established pathway databases for circRNAs. And although multiple mechanisms of circRNA function have been proposed, it remains largely unclear which circRNAs regulate which downstream mRNAs and through which mechanisms. Current approaches, such as circRNA–miRNA–mRNA network construction [59], provide insights but are primarily bioinformatic and motif-based, lacking experimental validation. Many circRNAs in our dataset have been studied in other contexts, particularly cancer, where they influence cell proliferation and signaling, but these findings are difficult to extrapolate to neuropsychiatric settings. For example, circSLTM, a TWAS prioritized circRNA shared across BIP, SCZ and AD, has been implicated in at-least two circRNA-miRNA-mRNA regulatory axes. One involves miR-421/HMGB2 axis associated with reduced neuronal conversion [60, 61]. Another involves the miR-515-5p/VAPB axis, which has been associated with attenuation of inflammation through NF-κB pathway [62]. Although the implications in psychiatry remain uncertain, encouraging examples are beginning to emerge. For instance, circHomer1 linked to regulating synaptic plasticity and cognition [29]. Similarly, circIgfbp2 connects neural plasticity and anxiety through mitochondrial dysfunction and oxidative stress-induced synapse impairment after traumatic brain injury [63]. Further, synaptically enriched circRNAs identified in rat hippocampal neurons were shown to regulate excitatory synapse formation via miRNA-mediated mechanisms [58], while in another study mouse midbrain dopamine neuron–specific circRNAs were linked to neuronal morphology and migration [64]. These examples highlight the potential of circRNAs in neuropsychiatric research and underscore the need for deeper mechanistic studies. We hope our work provides candidate circRNAs for future investigations, offering an opportunity to advance understanding of their roles in brain function and disease.

Our findings should be interpreted in the context of several technical and biological limitations. Low representation of circRNAs in ribosome-depleted RNA-seq libraries, due to their relative lower abundance compared to linear RNAs, could cause both false positives and false negatives, even with rigorous quality control, because of inherent noise. Short-read sequencing further complicates accurate circRNA quantification: when back-splice junctions are missed, reads originating from circular RNAs can be incorrectly aligned to linear transcripts. This misalignment inflates linear RNA counts because the pipeline interprets these reads as coming from linear isoforms rather than circular ones, leading to inaccurate representation of both RNA species. To mitigate this, our analysis incorporated both linear-gene and linear-junction measures, but advanced approaches such as long-read sequencing and improved profiling technologies will be essential for precise circRNA characterization. Integrating miRNA data from the same samples would strengthen circRNA–miRNA network construction, as current methods rely on external datasets and motif-based predictions. Beyond computational networks, many functional impacts of circRNAs remain unknown and uncaptured in bulk transcriptomic studies. Cell-type specificity, with single-cell approaches, could provide higher-resolution insights. To conclude, our work identifies SCZ- and BIP-associated circRNA signatures and supports circRNAs as a distinct regulatory molecular layer linked to psychiatric disease biology. Future progress will depend on consolidating larger datasets and employing carefully designed RNA profiling strategies, complemented by in vivo experiments to explore mechanistic hypotheses.

## Author Contribution

ND conceived the study. AJ and ND acquired funding. AJ and ND designed the analytical pipelines. AJ and MM implemented the computational pipelines. AJ and ND interpreted the results. AJ drafted the manuscript. MM contributed to writing the Methods and figure legends. All authors reviewed, edited, and approved the final manuscript.

## Acknowledgements

We thank members of the Daskalakis Lab for helpful discussions regarding the presentation and interpretation of our findings. We thank Shivachetan Ulavi for assistance with running the find_circ pipeline to generate circRNA counts, Vijetha Balakundi for imputing circGReX in the UK Biobank dataset, and Artemis Iatrou for preliminary pathway enrichment analyses. We also thank the PsychENCODE Consortium for providing access to the datasets used in this study, and the families of brain donors whose generous participation made this work possible.

## Methods

The PEC Phase 1 DLPFC dataset, comprising five studies (CMC, CMC-HBCC, LIBD, BrainGVEX, and UCLA-ASD), was downloaded from Synapse (node: syn8466658)[28, 65]. Only samples with rRNA-depleted libraries were retained for downstream analysis, as circRNAs are absent in poly-A libraries (N=1022; Supplementary Table 1-2). PEC genotypes were generated and imputed as previously reported [28]. Multi-allelic SNPs were removed, and genotype dosages were computed on continuous scale from 0 (homozygous reference) to 2 (homozygous alternate), with 1 representing heterozygous genotypes. We calculate ancestry proportions for each subject by projecting their genotype data onto principal components derived from known continental level ancestry markers. For this we used global ancestry pipeline (https://github.com/nievergeltlab/global_ancestry. PEC tissue preparation, RNA extraction, cDNA library preparation, and sequencing procedures were reported in We adopted the GTEx RNA-seq processing pipeline [66] for linear quantification and find_circ2 [67] for circRNA quantification. To name the 10, 936 circRNAs identified in our study, we followed the system proposed by Chen et al. [68]. Normalization and processing fo RNAseq counts was performed using DESeq2.

We roughly follow network analysis steps of Dube et. Al. [26] to build co-expression network. We used circNetVis [59] to explore circRNA–miRNA–mRNA interactions, applying default parameters. circRNA–miRNA interactions were predicted using three algorithms: TargetScan, RNAhybrid, and miRanda [69–71]. *metascape was used to perform DisGenNet and GO enrichments* [72, 73]. RNA-seq profiles of 382 neurotypical subjects from European ancestry were used for GReX model training. We adopted metaXcan database construction methodology to make our prediction model output compatible with the metaXcan software family. We used GCTA pipeline to estimate cis-heritability scores [73]. We performed TWAS using S-PrediXcan [74] on GWAS summary statistics from **a)** the *primary* SCZ GWAS (cases = 71,554; controls = 97,863) [75], which included participants of European, Admixed African, East Asian, and Latinx ancestries **b)** BIP GWAS (cases = 41917; controls = 371549) [76], and c) AD GWAS (cases = 111,326; controls = 677,663) [77]. GTAs (Gene Trait Associations) were deemed significant (bonf-GTAs) at a Bonferroni threshold of 0.05. This work was conducted under approval from the UK Biobank (UKBB application: 51979). circGReX models were used to impute genetically regulated circRNA expression (icirc) in ∼500,000 UK Biobank participants using MetaXcan. Associations between icirc and imaging or behavioral phenotypes were tested using linear regression (y∼icirc+covariates). The covariates included age, sex, genetic PCs, and imaging technical variables to account for confounding factors. We tested co-expression modules containing FDR-circDEGs for enrichment of TWAS-prioritized genes as reposrted in [37]. For this analysis, we defined four TWAS categories: i) SCZ circRNA TWAS associations identified in this study, ii) SCZ TWAS-prioritized linear genes reported in the literature, iii) psychiatric disorder TWAS-prioritized genes reported in the literature, including SCZ, depression, BIP, post-traumatic stress disorder, and autism spectrum disorder, and iv) neurodegenerative disorder TWAS-prioritized genes reported in the literature, including AD, frontotemporal dementia, Parkinson’s disease, and Lewy body dementia.

## Reference

1. Gratten, J., et al., Large-scale genomics unveils the genetic architecture of psychiatric disorders. Nat Neurosci, 2014. 17(6): p. 782–90.

2. Andreassen, O.A., et al., New insights from the last decade of research in psychiatric genetics: discoveries, challenges and clinical implications. World Psychiatry, 2023. 22(1): p. 4–24.

3. Trubetskoy, V., et al., Mapping genomic loci implicates genes and synaptic biology in schizophrenia. Nature, 2022. 604(7906): p. 502–508.

4. Mullins, N., et al., Genome-wide association study of more than 40,000 bipolar disorder cases provides new insights into the underlying biology. Nat Genet, 2021. 53(6): p. 817–829.

5. Gamazon, E.R., et al., A gene-based association method for mapping traits using reference transcriptome data. Nat Genet, 2015. 47(9): p. 1091–8.

6. Gusev, A., et al., Integrative approaches for large-scale transcriptome-wide association studies. Nat Genet, 2016. 48(3): p. 245–52.

7. Bhattacharya, A., et al., Isoform-level transcriptome-wide association uncovers genetic risk mechanisms for neuropsychiatric disorders in the human brain. Nature Genetics, 2023. 55(12): p. 2117–2128.

8. Gusev, A., et al., Transcriptome-wide association study of schizophrenia and chromatin activity yields mechanistic disease insights. Nature Genetics, 2018. 50(4): p. 538–548.

9. Gandal, M.J., et al., Shared molecular neuropathology across major psychiatric disorders parallels polygenic overlap. Science, 2018. 359(6376): p. 693–697.

10. Fromer, M., et al., Gene expression elucidates functional impact of polygenic risk for schizophrenia. Nature Neuroscience, 2016. 19(11): p. 1442–1453.

11. JaXe, A.E., et al., Developmental and genetic regulation of the human cortex transcriptome illuminate schizophrenia pathogenesis. Nat Neurosci, 2018. 21(8): p. 1117–1125.

12. Gandal, M.J., et al., Transcriptome-wide isoform-level dysregulation in ASD, schizophrenia, and bipolar disorder. Science, 2018. 362(6420).

13. Vattathil, S.M., et al., Mapping the microRNA landscape in the older adult brain and its genetic contribution to neuropsychiatric conditions. Nat Aging, 2025. 5(2): p. 306–319.

14. Nersisyan, S., et al., Several novel classes of small regulatory RNAs show widespread changes in schizophrenia and bipolar disorder and extensive linkages to critical brain processes. Translational Psychiatry, 2026. 16(1): p. 72.

15. Casey, C., J.F. Fullard, and R.D. Sleator, Unravelling the genetic basis of Schizophrenia. Gene, 2024. 902: p. 148198.

16. Greer, T.L., Circular RNAs as putative biomarkers for depression diagnosis and treatment. EBioMedicine, 2021. 68: p. 103362.

17. Khan, I.M., et al., Circular RNA Expression and Regulation Profiling in Testicular Tissues of Immature and Mature Wandong Cattle (Bos taurus). Front Genet, 2021. 12: p. 685541.

18. Gu, A., et al., Functions of Circular RNA in Human Diseases and Illnesses. Noncoding RNA, 2023. 9(4).

19. Rybak-Wolf, A., et al., Circular RNAs in the Mammalian Brain Are Highly Abundant, Conserved, and Dynamically Expressed. Mol Cell, 2015. 58(5): p. 870–85.

20. You, X., et al., Neural circular RNAs are derived from synaptic genes and regulated by development and plasticity. Nature Neuroscience, 2015. 18(4): p. 603–610.

21. Mahmoudi, E., M.J. Green, and M.J. Cairns, Dysregulation of circRNA expression in the peripheral blood of individuals with schizophrenia and bipolar disorder. J Mol Med (Berl), 2021. 99(7): p. 981–991.

22. Mahmoudi, E., et al., Circular RNA biogenesis is decreased in postmortem cortical gray matter in schizophrenia and may alter the bioavailability of associated miRNA. Neuropsychopharmacology, 2019. 44(6): p. 1043–1054.

23. Liu, Z., et al., Detection of circular RNA expression and related quantitative trait loci in the human dorsolateral prefrontal cortex. Genome Biol, 2019. 20(1): p. 99.

24. Mai, T.-L., et al., Trans-genetic eWects of circular RNA expression quantitative trait loci and potential causal mechanisms in autism. Molecular Psychiatry, 2022. 27(11): p. 4695–4706.

25. Chen, Y.J., et al., Genome-wide, integrative analysis of circular RNA dysregulation and the corresponding circular RNA-microRNA-mRNA regulatory axes in autism. Genome Res, 2020. 30(3): p. 375–391.

26. Dube, U., et al., An atlas of cortical circular RNA expression in Alzheimer disease brains demonstrates clinical and pathological associations. Nat Neurosci, 2019. 22(11): p. 1903–1912.

27. Lo, I., et al., Linking the association between circRNAs and Alzheimer’s disease progression by multi-tissue circular RNA characterization. RNA Biol, 2020. 17(12): p. 1789–1797.

28. Wang, D., et al., Comprehensive functional genomic resource and integrative model for the human brain. Science, 2018. 362(6420).

29. Zimmerman, A.J., et al., A psychiatric disease-related circular RNA controls synaptic gene expression and cognition. Molecular Psychiatry, 2020. 25(11): p. 2712–2727.

30. Song, W.M. and B. Zhang, Multiscale Embedded Gene Co-expression Network Analysis. PLoS Comput Biol, 2015. 11(11): p. e1004574.

31. Langfelder, P. and S. Horvath, WGCNA: an R package for weighted correlation network analysis. BMC Bioinformatics, 2008. 9(1): p. 559.

32. Hochman, K.M. and R.R. Lewine, Age of menarche and schizophrenia onset in women. Schizophrenia Research, 2004. 69(2): p. 183–188.

33. Ashwal-Fluss, R., et al., circRNA biogenesis competes with pre-mRNA splicing. Mol Cell, 2014. 56(1): p. 55–66.

34. Grotzinger, A.D., et al., Mapping the genetic landscape across 14 psychiatric disorders. Nature, 2026. 649(8096): p. 406–415.

35. Wingo, T.S., et al., Shared mechanisms across the major psychiatric and neurodegenerative diseases. Nature Communications, 2022. 13(1): p. 4314.

36. Yu, A.W., J.D. Peery, and H. Won, Limited Association between Schizophrenia Genetic Risk Factors and Transcriptomic Features. Genes, 2021. 12(7): p. 1062.

37. Gao, H., et al., TWAS atlas 2.0: an updated data resource for transcriptome-wide association studies. Nucleic Acids Res, 2026. 54(D1): p. D1321–D1330.

38. Gerring, Z.F., et al., An analysis of genetically regulated gene expression and the role of co-expression networks across 16 psychiatric and substance use phenotypes. European Journal of Human Genetics, 2022. 30(5): p. 560–566.

39. Soo, C.C., et al., Genome-wide association study of population-standardised cognitive performance phenotypes in a rural South African community. Communications Biology, 2023. 6(1): p. 328.

40. Zeng, L., et al., A Single-Nucleus Transcriptome-Wide Association Study Implicates Novel Genes in Depression Pathogenesis. Biol Psychiatry, 2024. 96(1): p. 34–43.

41. Wang, F., et al., Integration of multiple-omics data to reveal the shared genetic architecture of educational attainment, intelligence, cognitive performance, and Alzheimer’s disease. Frontiers in Genetics, 2023. **Volume 14 -** 2023.

42. Arizaca Maquera, K.A., et al., Alzheimer’s disease pathogenetic progression is associated with changes in regulated retained introns and editing of circular RNAs. Front Mol Neurosci, 2023. 16: p. 1141079.

43. Memczak, S., et al., Circular RNAs are a large class of animal RNAs with regulatory potency. Nature, 2013. 495(7441): p. 333–338.

44. Nguyen, T.-H., H.-N. Nguyen, and T.-N. Vu, CircNetVis: an interactive web application for visualizing interaction networks of circular RNAs. BMC Bioinformatics, 2024. 25.

45. Elliott, L.T., et al., Genome-wide association studies of brain imaging phenotypes in UK Biobank. Nature, 2018. 562(7726): p. 210–216.

46. Hanan, M., et al., A Parkinson’s disease CircRNAs Resource reveals a link between circSLC8A1 and oxidative stress. EMBO Mol Med, 2020. 12(9): p. e11942.

47. Yi, G., et al., circKLHL24 Blocks Breast Cancer Development by Regulating the miR-1204/ALX4 Network. Cancer Biother Radiopharm, 2022. 37(8): p. 684–696.

48. Kayserili, H., et al., ALX4 dysfunction disrupts craniofacial and epidermal development. Human Molecular Genetics, 2009. 18(22): p. 4357–4366.

49. Wu, W., P. Ji, and F. Zhao, CircAtlas: an integrated resource of one million highly accurate circular RNAs from 1070 vertebrate transcriptomes. Genome Biol, 2020. 21(1): p. 101.

50. Dong, X., et al., Circular RNAs in the human brain are tailored to neuron identity and neuropsychiatric disease. Nature Communications, 2023. 14(1): p. 5327.

51. Huang, W., et al., TransCirc: an interactive database for translatable circular RNAs based on multi-omics evidence. Nucleic Acids Research, 2020. 49(D1): p. D236–D242.

52. Li, X., L. Yang, and L.-L. Chen, The Biogenesis, Functions, and Challenges of Circular RNAs. Molecular Cell, 2018. 71(3): p. 428–442.

53. Liang, D., et al., The Output of Protein-Coding Genes Shifts to Circular RNAs When the Pre-mRNA Processing Machinery Is Limiting. Mol Cell, 2017. 68(5): p. 940–954 e3.

54. Liu, H., et al., Identification of genetic architecture shared between schizophrenia and Alzheimer’s disease. Translational Psychiatry, 2025. 15(1): p. 150.

55. Gerring, Z.F., E.R. Gamazon, and E.M. Derks, A gene co-expression network-based analysis of multiple brain tissues reveals novel genes and molecular pathways underlying major depression. PLoS Genet, 2019. 15(7): p. e1008245.

56. Borcuk, C., et al., Network-wide risk convergence in gene co-expression identifies reproducible genetic hubs of schizophrenia risk. Neuron, 2024. 112(21): p. 3551–3566.e6.

57. GeoXroy, P.A., et al., Combination therapy for manic phases: a critical review of a common practice. CNS Neurosci Ther, 2012. 18(12): p. 957–64.

58. Kelly, D., et al., A functional screen uncovers circular RNAs regulating excitatory synaptogenesis in hippocampal neurons. Nature Communications, 2025. 16(1): p. 3040.

59. Nguyen, T.-H., H.-N. Nguyen, and T.N. Vu, CircNetVis: an interactive web application for visualizing interaction networks of circular RNAs. BMC Bioinformatics, 2024. 25(1): p. 31.

60. Zhang, H., et al., Circular RNA SLTM as a miR-421-competing endogenous RNA to mediate HMGB2 expression stimulates apoptosis and inflammation in arthritic chondrocytes. J Biochem Mol Toxicol, 2023. 37(5): p. e23306.

61. 61. Lepko, T., The role of chromatin associated protein HMGB2 in setting up permissive chromatin states for direct glia to neuron conversion. 2018.

62. Chen, R., et al., circSLTM knockdown attenuates chondrocyte inflammation, apoptosis and ECM degradation in osteoarthritis by regulating the miR-515-5p/VAPB axis. Int Immunopharmacol, 2024. 138: p. 112435.

63. Du, M., et al., A novel circular RNA, circIgfbp2, links neural plasticity and anxiety through targeting mitochondrial dysfunction and oxidative stress-induced synapse dysfunction after traumatic brain injury. Mol Psychiatry, 2022. 27(11): p. 4575–4589.

64. Rybiczka-Tešulov, M., et al., Circular RNAs regulate neuron size and migration of midbrain dopamine neurons during development. Nature Communications, 2024. 15(1): p. 6773.

65. Akbarian, S., et al., The PsychENCODE project. Nature Neuroscience, 2015. 18(12): p. 1707–1712.

66. Consortium, G.T., The GTEx Consortium atlas of genetic regulatory eWects across human tissues. Science, 2020. 369(6509): p. 1318–1330.

67. Memczak, S., et al., Circular RNAs are a large class of animal RNAs with regulatory potency. Nature, 2013. 495(7441): p. 333–8.

68. Chen, L.L., et al., A guide to naming eukaryotic circular RNAs. Nat Cell Biol, 2023. 25(1): p. 1–5.

69. McGeary, S.E., et al., The biochemical basis of microRNA targeting eWicacy. Science, 2019. 366(6472).

70. Krüger, J. and M. Rehmsmeier, RNAhybrid: microRNA target prediction easy, fast and flexible. Nucleic Acids Res, 2006. 34(Web Server issue): p. W451–4.

71. Enright, A.J., et al., MicroRNA targets in Drosophila. Genome Biol, 2003. 5(1): p. R1.

72. Piñero, J., et al., DisGeNET: a comprehensive platform integrating information on human disease-associated genes and variants. Nucleic Acids Research, 2017. 45(D1): p. D833–D839.

73. Zhou, Y., et al., Metascape provides a biologist-oriented resource for the analysis of systems-level datasets. Nat Commun, 2019. 10(1): p. 1523.

74. Barbeira, A.N., et al., Exploring the phenotypic consequences of tissue specific gene expression variation inferred from GWAS summary statistics. Nat Commun, 2018. 9(1): p. 1825.

75. Trubetskoy, V., et al., Mapping genomic loci implicates genes and synaptic biology in schizophrenia. Nature, 2022. 604(7906): p. 502–508.

76. Mullins, N., et al., Genome-wide association study of more than 40,000 bipolar disorder cases provides new insights into the underlying biology. Nature Genetics, 2021. 53(6): p. 817–829.

77. Bellenguez, C., et al., New insights into the genetic etiology of Alzheimer’s disease and related dementias. Nature Genetics, 2022. 54(4): p. 412–436.

